# Horizontal gene transfer rivals gene duplication as a source of anti-parasitoid immune innovation in the Drosophilidae

**DOI:** 10.64898/2026.09.15.751804

**Authors:** Rebecca L. Tarnopol, Ryan L. Wang, Bernard Y. Kim, Noah K. Whiteman

**Affiliations:** Department of Molecular & Cell Biology, University of California, Berkeley, CA 94720; Department of Microbiology & Immunology, University of California, San Francisco, CA 94158; Department of Ecology & Evolutionary Biology, Princeton University, Princeton, NJ 08544; Department of Integrative Biology, University of California, Berkeley, CA 94720

**Keywords:** Innate immunity, Drosophila, horizontal gene transfer, PPO, parasitism

## Abstract

Macroparasites are among the most important agents of natural selection in their host populations, but anti-macroparasite immunity is poorly understood. Vinegar flies (Drosophilidae) and the parasitoid wasps that infect them are emerging models to study how animals defend against macroparasite attack. The canonical anti-parasitoid immune mechanism in insects is melanotic encapsulation, which involves cell-mediated encapsulation coupled with prophenoloxidase (PPO)-mediated melanization of parasitoid embryos. Recently, we discovered the horizontal transfer (HGT) of a bacterially-derived humoral anti-parasitoid effector, Cytolethal distending toxin B (CdtB), across insects, including four drosophilid lineages. Here, we assessed the prevalence of these two anti-parasitoid immune mechanisms in 406 drosophilid and four outgroup species. We found that melanotic encapsulation was relatively uncommon among species with known anti-parasitoid immune responses. While *PPO* duplications were found in 88 species, the most salient *PPO* gene underlying melanotic encapsulation, *PPO3*, was restricted to *Drosophila melanogaster* and its close relatives. *PPO* genes were present in lower copy number per genome in the Drosophilidae than in outgroup lineages and evolved slowly. We found *cdtB* in the genomes of 93 species and estimated at least 11 independent *cdtB* gains across the Drosophilidae as early as ∼44 mya and as recently as ∼6 mya. *cdtB* acquisition was subsequently associated with higher net diversification rates in some clades. We conclude that humoral immune effectors may play a more important role than previously appreciated in anti-parasitoid immunity in insects and that an immune innovation arising repeatedly from HGT is potentially associated with the evolutionary success of these animals.

**Significance Statement:** Insects have evolved robust immune strategies to overcome parasitoid wasp challenge. The prevailing mechanism involves encapsulation of immature parasitoids using melanin-producing blood cells dependent on duplicated prophenoloxidase (*PPO*) genes. Another anti-parasitoid response evolved through the horizontal gene transfer (HGT) of a toxin-encoding gene (*cdtB*) from endosymbionts to the genomes of insects. We show the number of species encoding *PPO* duplications and species encoding *cdtB* were similar in the Drosophilidae. HGT, but not *PPO* duplication, was associated with higher rates of diversification within some fly lineages, indicating that the HGT events or an associated trait may confer a fitness advantage. These results suggest that anti-parasitoid immune mechanisms that act independently of melanotic encapsulation responses are effective and pervasive in the Drosophilidae.

## Introduction

Long-lived macroparasitic animals, including blood-feeding helminths, have played a major role in shaping the evolution of the immune system (1). However, experimental studies that aim to identify the molecular basis of anti-parasite defenses have been difficult owing to the specialized nature of these parasites and their experimental intractability (2, 3). *Drosophila melanogaster* and its parasitoid wasps are emerging as a useful model to address this gap. In natural populations of the vinegar fly *D. melanogaster*, wasp-induced mortality can rise to ∼90% annually (4). Parasitoid wasp larval life histories share important features with those of blood-feeding helminths: Both must survive in the circulatory system or hemocoel of their hosts, where they not only feed on blood or hemolymph, but also must endure the onslaught of the host immune system.

Despite their importance, little is known about the immune mechanisms used by insect hosts to defend against parasitoid wasps relative to the well-studied antimicrobial responses (5). The best understood anti-parasitoid immune response is that of *D. melanogaster*, which has a sophisticated cellular immune response in fly larvae against parasitoid wasps (6). Following parasitoid attack, a signaling cascade mediated primarily by the Toll and JAK-STAT pathways leads to the proliferation and differentiation of circulating blood cells (hemocytes) into specialized, wasp-encapsulating cells called lamellocytes (7–9). Lamellocytes then bind to the wasp embryo where they, in concert with other circulating hemocytes called crystal cells, neutralize the developing embryo through melanization reactions catalyzed by prophenoloxidases (PPOs). The primary PPO involved in this reaction, PPO3, arose from a gene duplication event of *PPO2*, with which *PPO3* shares its exon/intron structure (10, 11). This energetically expensive immune reaction is known as melanotic encapsulation and is hypothesized to be the primary immune mechanism by which other insects defend themselves against parasitoid wasp attack (12). Analogous pathways have been identified in model lepidopteran species and in other arthropods (13–15).

However, recent cell biology and genomic studies across the Drosophilidae have revealed that lamellocytes and *PPO3* are only found in *D. melanogaster* and its close relatives (10, 11, 16). Although anti-parasitoid immune responses have only been studied in the laboratory in a small fraction of extant species, only 18/51 (35.3%) performed melanotic encapsulation, the majority of which were close relatives of *D. melanogaster* (Figure 1A, Supplementary File S1). Most species assayed instead used multiple mechanisms for parasitoid neutralization, including 29/51 (56.8%) species with mechanisms independent of both encapsulation and melanization (Figure 1A, Supplementary File S1). Thus, even with sparse sampling, the anti-parasitoid immune mechanisms in drosophilids varied widely and were not dominated by cellular encapsulation and/or melanization.

**Figure 1.**
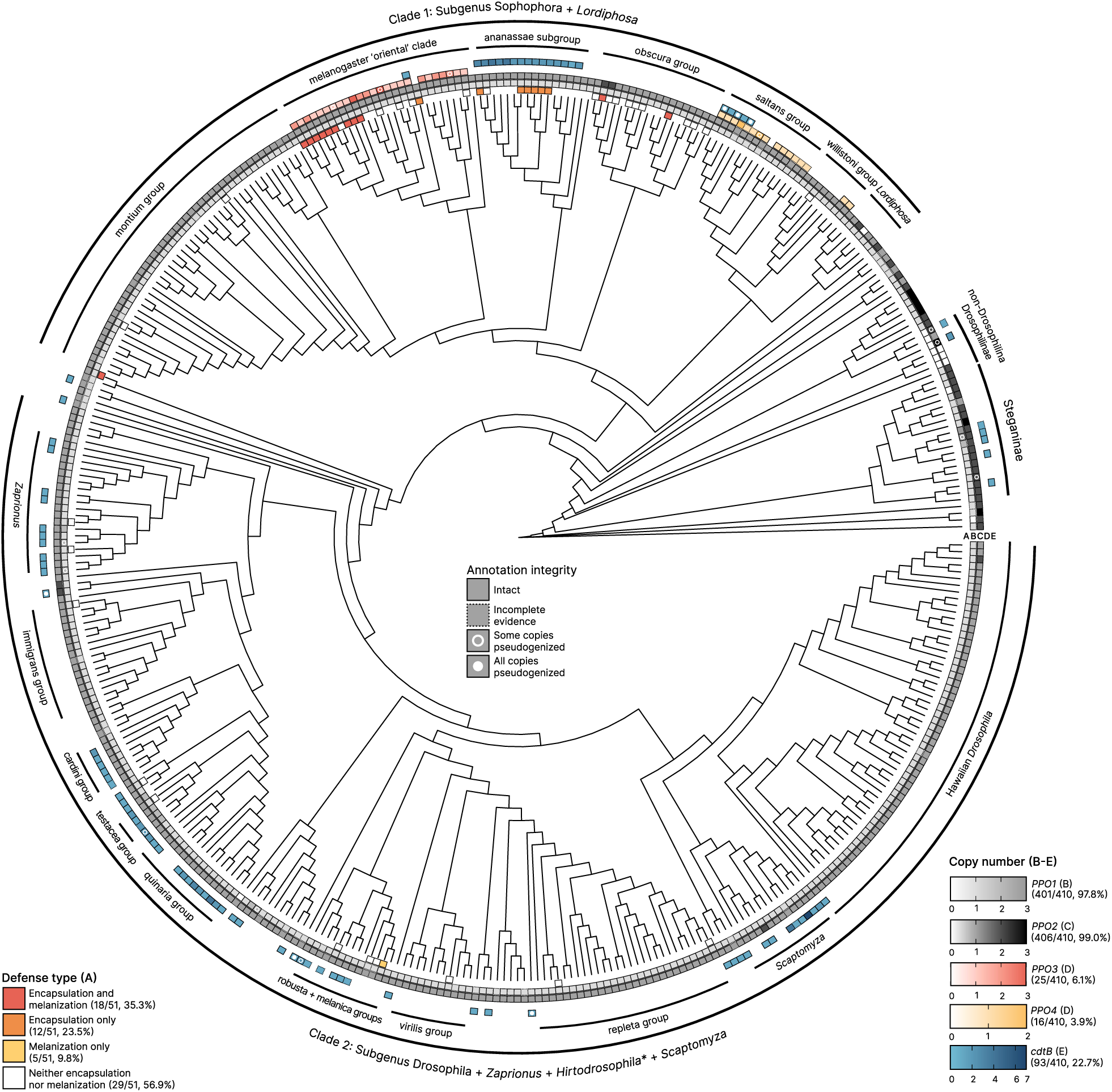
Melanization phenotypes are uncommon and associated *prophenoloxidase* duplications are about equal in prevalence to horizontally transferred humoral immune effectors in Drosophilidae. A maximum likelihood species tree comprising 406 drosophilid species and four outgroup taxa were surveyed for defense phenotypes, *PPO* gene content, and *cdtB* presence. Characters mapped from the inner ring outward: Defense type (A), *PPO1* copy number (B), *PPO2* copy number (C), *PPO3* and *PPO4* copy number (D), and *cdtB* copy number (E). Major taxa are labeled on the outermost rings. Clade 2 includes all *Hirtodrosophila* species sampled except for *H. duncani*, which is found in Clade 1. Branch lengths are equalized for clarity. Detailed view of the phylogeny may be viewed at https://trnpl.github.io/antiparasitoid/.

While the cellular arm of immunity remains an essential feature of the anti-parasitoid immune response in *D. melanogaster* and other arthropods, the paucity of this mechanism in many drosophilids indicates that alternative anti-parasitoid mechanisms have evolved across this diverse family. Indeed, recently we identified an anti-parasitoid response involving toxic humoral factors, similar in principle to the toxic humoral peptides deployed against fungi, oomycetes, and bacteria that led to the discovery of the role of the Toll pathway in mediating anti-microbial immunity in animals (5). Cytolethal distending toxin B (CdtB) is an apoptosis-inducing DNAse I homolog that was horizontally transferred from defensive endosymbiotic bacteria and their phages to the nuclear genomes of insects across several orders, including at least four lineages of drosophilid flies (17–22). In *Drosophila*, CdtB functions in the humoral immune system, constituting a novel anti-parasitoid response that acts independently of cellular encapsulation or melanization responses. Functional genetic studies found that CdtB homologs are both necessary for full anti-parasitoid immune responses in *D. ananassae*, where they are natively encoded, and sufficient to drive a robust anti-parasitoid phenotype when expressed heterologously in the *D. melanogaster* fat body (21, 23). These studies demonstrated that horizontal gene transfer (HGT) can confer rapid adaptation to parasitoid wasp pressure by introducing novel anti-parasitoid effectors to insect immune systems.

Genomic resources are now available for 400+ species of the Drosophilidae, allowing for phylogenomic approaches to be used to illuminate the prevalence of each of these responses across the phylogenetic diversity of this model insect family. Here, we used these rich genomic resources to dissect the evolution of two gene families encoding important anti-parasitoid factors: Prophenoloxidases (PPOs), responsible for parasitoid melanization in the canonical cellular anti-parasitoid defense mechanism (Figure 1B-D), and Cytolethal distending toxin subunit B (CdtB) proteins (Figure 1E), responsible for the toxic, humoral anti-parasitoid defense mechanism.

## Results

### Prophenoloxidases evolve slowly in the Drosophilidae

Prophenoloxidases are type-III copper-binding enzymes that catalyze melanization reactions. These reactions produce noxious superoxide species hypothesized to underlie the parasitoid killing mechanism during melanotic encapsulation (24). *D. melanogaster* encodes three prophenoloxidase genes with overlapping but non-redundant functions. *PPO1* and *PPO2* are expressed in crystal cells, where they are involved in wound healing and anti-microbial responses against Gram-positive bacteria and fungi (25). PPO2 also has additional roles in larval physiology, including oxygen transport (26). *PPO3* is a *PPO2* paralog, and while both PPO2 and PPO3 contribute to melanizing encapsulation in *D. melanogaster*, PPO3 is the major anti-parasitoid prophenoloxidase and is expressed in parasitoid-encapsulating lamellocytes (10, 11, 27).

We manually curated annotations for all *PPO* genes in 406 drosophilid and four outgroup species with completely sequenced genomes (see Materials & Methods, SI Methods). We then created an interactive web browser to visualize these data as well as *cdtB* annotation data and other species phylogeny-wide analyses conducted in this study (see Data & Materials Availability). We assigned *PPO* orthology based on shared syntenic location with the paralogs in *D. melanogaster* and then validated their identities by constructing a nucleotide phylogeny. We found that *PPO1* and *PPO2* were highly conserved in the Drosophilidae (Figure 1B-C), and with the exception of *Scaptomyza polygonia PPO1* and *Leucophenga* spp. *PPO2*, were encoded in the same locus across all screened species (Figure 2A). Tandem duplications of *PPO2* were common in Steganinae species and other non-*Drosophila* drosophilids. Consistent with previous reports, *PPO3* was restricted to *D. melanogaster* and its close relatives, comprising 25 species (6.2%) in the ‘oriental’ clade of the *melanogaster* species group. We also found a new *PPO2-*like paralog encoded by species in the *willistoni* and *saltans* groups, which we named *PPO4* given its monophyly and high divergence from the other *PPO*s (Figure 2B, Figure S1). In total, 88/406 (21.7%) drosophilid species encoded >2 *PPO* copies. While most species capable of melanotic encapsulation have *PPO* duplications, seven species melanized parasitoids while encoding only *PPO1* and *PPO2* (Figure 1A, Supplementary File S1), showing that melanization responses can arise without *PPO* duplications.

**Figure 2.**
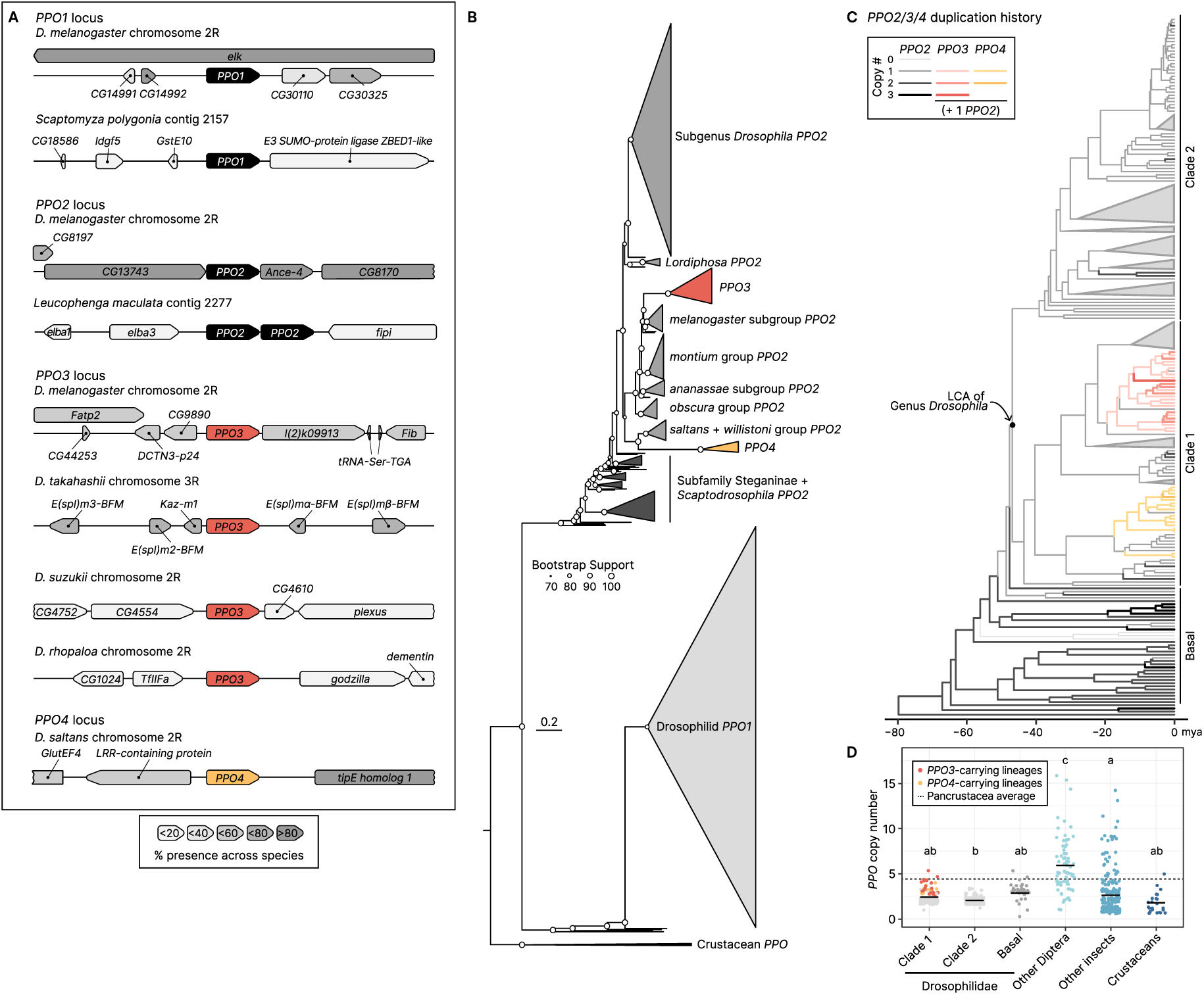
PPO genes evolve relatively slowly in the Drosophilidae. (A) Cartoons of syntenic loci for each of the PPO paralogs recovered in Drosophilidae. Neighboring genes are shaded per their relative prevalence in all taxa that encode each paralog. Scaling of neighboring genes is relative to each *PPO*, but does not follow a fixed scale. (B) A maximum likelihood nucleotide phylogeny (best fit model: GTR+F+I+R9) of *PPO* paralogs. The tree was rooted at the clade comprising crustacean *PPO* homologs. Scale bar = substitutions/site. (C) Timing of *PPO2/3/4* duplication events across the Drosophilidae as inferred from CAFE5. Collapsed clades and are scaled to 20% size relative to species richness. Branch thickness indicates copy number, and *PPO3-* and *PPO4-*encoding lineages are highlighted in pink and yellow, respectively. (D) *PPO* copy number variation in the Drosophilidae and outgroup taxa in pancrustacea. Dashed line indicates the average *PPO* copy number across all orders in the pancrustacea. Letters indicate significant difference between groups as determined by ANOVA followed by Tukey’s test. A detailed view of the CAFE5 reconstruction for *PPO2*/*3*/*4* (as shown in 2C) and for *PPO1* can be found at https://trnpl.github.io/antiparasitoid/.

Similar to *PPO1* and *PPO2*, all *PPO4* genes in the *saltans* and *willistoni* species groups were in the same locus (Figure 2A). In contrast, *PPO3* was in four different syntenic locations (Figure 2A). *PPO3* was most commonly found embedded between *E(spl)* genes, a series of Notch repressors active during the earliest stages of prohemocyte differentiation (Figure 2A) (28). Nine species encoded *PPO3* in more than one of these locations, indicating rampant duplication and loss dynamics in the evolution of this paralog. At least partial loss of *PPO3* was found in two species: *D. ficusphila*, from which we were unable to recover any *PPO3* sequences despite high contiguity in all observed *PPO3* loci, and *D. sechellia*, which encodes a pseudogenized version of *PPO3* in its reference genome (10, 11). However, we discovered a segregating polymorphism for a functional *PPO3* allele in *D. sechellia* that was previously unknown (Figure 1D, Figure S2). Notably, these two species did not melanotically encapsulate parasitoids in the laboratory, serving as natural loss of function mutants that underscore the role of *PPO3* in mediating this immune response in this lineage (Figure 1A) (10, 29).

In the maximum likelihood nucleotide phylogeny, *PPO3* homologs formed a long branch stemming from *PPO2* orthologs encoded by *melanogaster* subgroup species, as did the *PPO4* homologs from *PPO2* orthologs encoded by *willistoni* and *saltans* group species (Figure 2B). These data, combined with CAFE5 ancestral state estimation (Figure 2C), indicate that these paralogs arose from recent *PPO2* gene duplication events in the last common ancestor of each of these clades ∼17 mya.

Gene duplication events are a major source of evolutionary novelty in animals (30–32). We next tested whether *PPO* family genes turned over rapidly or had signatures of diversifying selection that are associated with duplication events. Neither *PPO1* nor *PPO2/3/4* gene families experienced significant shifts in gene family size over the course of evolution in the Drosophilidae (P > 0.99), though *PPO2/3/4* turned over faster than *PPO1* (γ*_PPO2/3/4_* = 1.455 vs. γ*_PPO1_* = 0.545, Table S1). We found evidence for episodic positive diversifying selection for each paralog using BUSTED analyses, though average ⍵ was < 1 in all cases (Table S2). Branch site tests conducted in aBSREL revealed evidence for diversifying selection specifically on branches leading to clades of *PPO2* duplications, consistent with accelerated rates of evolution after an initial duplication event in the last common ancestor of a lineage, followed by purifying selection in branches leading to extant lineages (Table S2). Taken together, our results show that *PPO* homologs in the Drosophilidae have evolved relatively slowly after the initial *PPO2* duplication, suggesting melanization reactions are not frequently exapted for parasitoid defense in the family.

Strikingly, CAFE5 predicted two *PPO2* copies as the ancestral state with subsequent loss in the branch leading to the radiation of genus *Drosophila* (LCA of Clade 1 and Clade 2, Figure 2C). Most secondary *PPO2* duplication events (5/8, 62.5%) in this radiation occurred in species in the clade inclusive of subgenus *Sophophora* and genus *Lordiphosa* species (“Clade 1”). Because all outgroup species we screened had *PPO* duplications, we hypothesized that *Drosophila* spp. are anomalous in their *PPO* copy number compared to other dipteran lineages, which are reported to have extensive copy number expansions in *PPO* family genes (33). Indeed, drosophilids were *PPO-*poor compared to other dipteran lineages, which had an average of 5.3 *PPO* copies/species (Figure 2D, Table S3). Basal Drosophilidae had the highest average *PPO* copy number at 2.86 copies/species, while Clade 2 species had the lowest, encoding on average 2.05 copies/species (Figure 2, Table S3). Clade 1 species had slightly elevated *PPO* copy numbers at 2.42 copies/species, largely driven by expansions in *PPO3-* and *PPO4-*encoding lineages (Figure 2D, Table S3).

We further screened all publicly available annotated genome assemblies from insect and crustacean species for *PPO* genes to determine if *Drosophila* spp. had a similar *PPO* copy number relative to other pancrustacean lineages, as *PPO* expansions are thought to be rare in insect lineages outside of Diptera (see Materials & Methods) (33, 34). This search revealed that *PPO* gene family expansions were much more common among insects than previously appreciated, with insects and crustaceans encoding an average of 4.4 *PPO* copies/species (Figure 2D, Table S3).

### Humoral anti-parasitoid effector proteins were repeatedly acquired via HGT across the evolution of the Drosophilidae

Given the relatively low prevalence of melanotic encapsulation and *PPO* duplications in Drosophilidae, we next focused on the humoral anti-parasitoid response across the family. We focused on *cdtB* because laboratory studies have shown it is both necessary and sufficient for anti-parasitoid immunity in two *Drosophila* species (21, 23). We reasoned that the presence of *cdtB* could therefore be a useful marker of similar humoral anti-parasitoid responses in other drosophilid species, just as *PPO3* and potentially other *PPO* duplications are useful markers of melanotic encapsulation. Surprisingly, we identified *cdtB* in 93/406 (22.9%) species, nearly four times the number that encode *PPO3* and five more species than encode any *PPO* duplication (Figure 1E).

We previously identified *cdtB* acquisitions in *Scaptomyza* + *D. primaeva*, the *ananassae* subgroup, *D. biarmipes*, and *Zaprionus bogoriensis* (19, 21). In most of these cases, CdtB was encoded by an intronless, single copy gene. However, in the *ananassae* subgroup, *cdtB* had a three-exon structure (as did the copy in *D. biarmipes*), and there were additional *cdtB* copies fused to sequences encoding the receptor binding domain of a different toxin, Apoptosis Inducing Protein of 56 kilodaltons (AIP56), which were found in synteny in endosymbiotic APSE phage toxin cassettes (19). These initial findings suggested that while *cdtB* was rare among drosophilid species, multiple HGT events introduced the gene independently to the family either through bacteria and their phages to insects or insect-to-insect transfers.

Our comparative genomic screen of 410 species uncovered *cdtB* homologs in many new lineages, particularly in Clade 2 drosophilids and in the subfamily Steganinae. We did not find *cdtB* in any of the four outgroup species. Manual curation of *cdtB* annotations revealed cryptic introns in many homologs, which lengthened previously identified open reading frames in *Scaptomyza* (Figure 3A, Figure S3). We also identified *ananassae*-like three-exon *cdtB* in long-diverged species. Strikingly, splice junctions were conserved in all of the three-exon copies of *cdtB,* except for those in the *saltans* group (Figure 3A). *Colocasiomyia alocasiae* had single copy *cdtB* upstream of *cdtB::aip56* fusion genes that shared splice junctions with those in the *ananassae* subgroup (Figure 3A). We also discovered extensive copy number variation in *cdtB* homologs, with up to seven copies in some species (Figure 1E).

**Figure 3.**
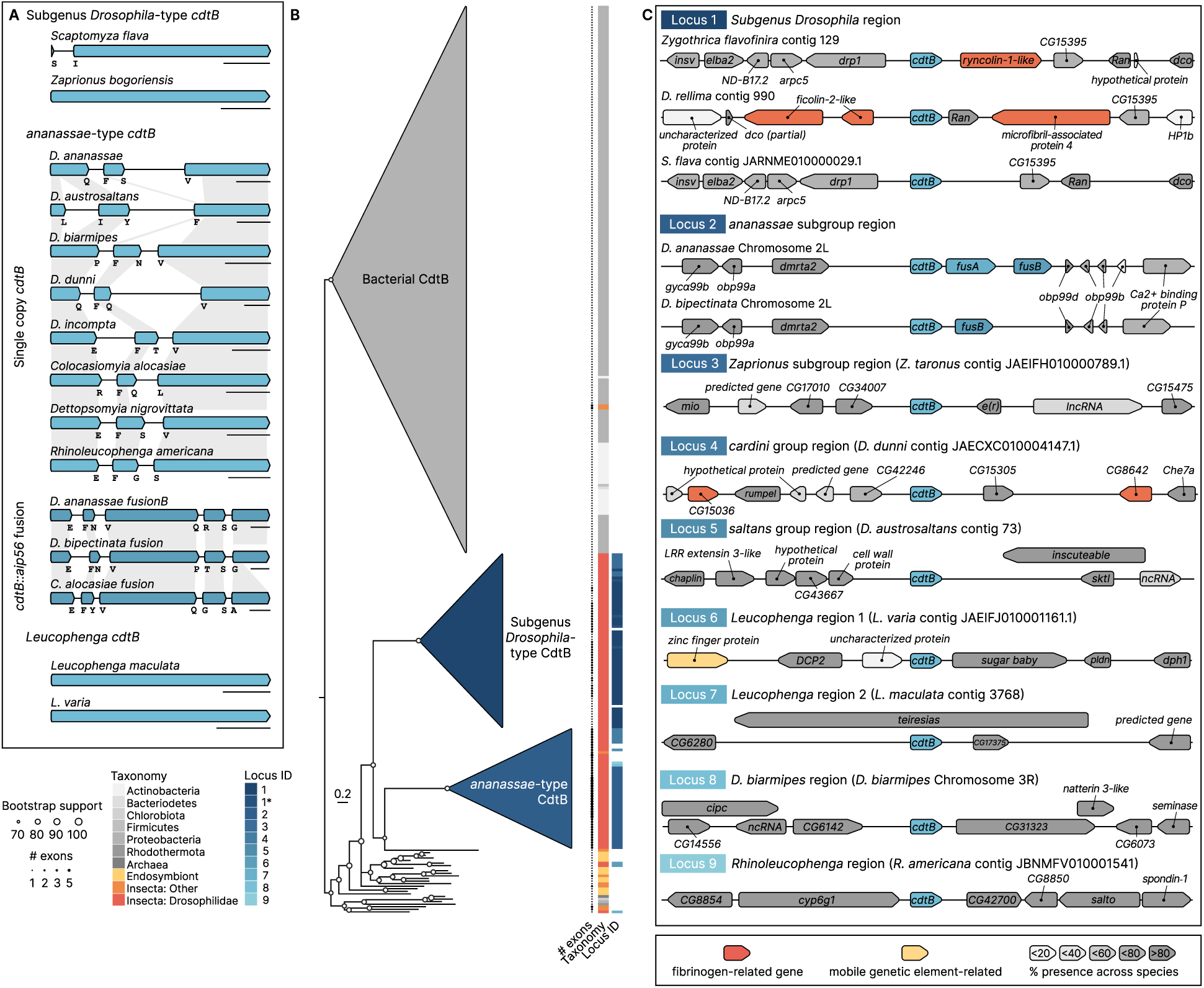
Drosophilid CdtBs form two major clades and are found in loci with other immune-related genes. (A) Gene models for four major drosophilid *cdtB* homologs identified in this study. Small letters indicate amino acids on either side of the splice junction, and grey boxes indicate homology. Scale bars = 200bp. (B) A maximum likelihood amino acid phylogeny (best fit model: WAG+F+R7) of CdtB homologs reveal two major clades of drosophilid CdtBs. The tree was rooted such that the clade of CdtB homologs from free-living bacteria was the outgroup. Exon count, taxonomy, and locus identity for drosophilid CdtBs are shown for each tip. Scale bar = substitutions/site. (C) Cartoons of unique syntenic loci identified across species. Neighboring genes are shaded per their relative prevalence in all taxa that encode *cdtB* at that locus. Fibrinogen-like genes and mobile genetic element-associated genes are shaded in red and yellow, respectively. Scaling of neighboring genes is relative to the size of *cdtB*, but does not follow a fixed scale.

The CdtB amino acid tree formed three major clades: CdtB homologs encoded by free-living bacteria, CdtB homologs from Clade 2 species (“subgenus *Drosophila*-type”), and insect-derived CdtB homologs that share a three-exon gene structure (“*ananassae*-type”, Figure 3B, Figure S4). CdtB homologs from insect endosymbionts (i.e. Ca. *Hamiltonella defensa*, *Arsenophonus* spp., *Wolbachia* spp., and their phages) were sister to insect homologs. *Leucophenga* spp. CdtB copies nested with those from endosymbionts and nematoceran Diptera, indicating distinct acquisition events from those that gave rise to other drosophilid *cdtB* genes (Figure 3A-B, Figure S4). The subgenus *Drosophila*-type CdtBs largely recapitulated the species tree topology, consistent with the hypothesis of a singular, ancient HGT event in the lineage’s last common ancestor. However, the *ananassae*-type CdtBs showed a more reticulated topology inconsistent with the species phylogeny (Figure S4). Notably, CdtB homologs from *D. incompta* and the *cardini* group formed a clade with homologs from *saltans* group species, which last shared a common ancestor before the split of subgenera *Sophophora* and *Drosophila* ∼46.9 mya. These copies were in turn sister to the CdtB homolog from the green peach aphid *Myzus persicae*, suggesting either inter-insect HGT events or ancient introgression between drosophilid species that shared ancestral geographical ranges.

To further estimate the number of independent *cdtB* acquisitions in the Drosophilidae, we characterized the microsyntenic loci proximal to *cdtB* in each species. Because prokaryote-to-animal HGT events followed by domestication are exceedingly rare, we scored shared microsynteny as a single ancient acquisition event in a common ancestor of the species. From sequences long enough to deduce gene neighborhoods, we observed nine putative ancestral chromosomal insertions for *cdtB* (Figure 3C). We found *cdtB* in the same locus in long-diverged species like *Zygothrica flavofinira* and *Scaptomyza flava,* further supporting a single acquisition event before the radiation of the subgenus *Drosophila* (Figure 3C). Additionally, we detected *cdtB* transposition after an initial gain in a number of lineages, including in *Zaprionus* (Figure 3C, Supplementary File S1).

Many *cdtB* homologs are in the proximity of other putative immune genes, suggesting these regions of the genome are primed for immune responses. *cdtB* homologs in subgenus *Drosophila* are frequently nested within arrays of fibrinogen- and ficolin-like genes, which are also present in sister taxa that do not encode *cdtB* (Figure 3C, Figure S5). Fibrinogens are important blood clotting factors and anti-helminth defenses in animals (35, 36), while ficolins act as pattern recognition receptors that promote opsonization of invading pathogens by activating the complement cascade (37). Fibrinogen-like genes were common in the subgenus *Drosophila* but largely absent in subgenus *Sophophora*, suggesting an important role for humoral effectors in the anti-parasitoid response in this lineage.

### *cdtB* acquisitions are both ancient and recurrent in the Drosophilidae

We next used ancestral state reconstruction (ASR) to predict the timing and frequency of *cdtB* acquisitions and losses throughout the drosophilid species phylogeny. To do this, we incorporated synteny and gene tree information to assign 12 distinct character states comprising 11 independent acquisition events and one transposition event to generate a total evidence model for ASR (see Materials & Methods). This model predicted on average 15 gains and 43 losses (Figure 4A, Table S4). The likely overestimation is due in part to uncertainties with the species tree topology, particularly in clades undergoing rapid diversification. This is evident in the Hawaiian drosophilid *D. primaeva* (which is sister to the picture-winged clade), in which *cdtB* was predicted as a secondary gain in both models despite sharing microsynteny with other subgenus *Drosophila cdtB* homologs, suggesting incomplete lineage sorting or an ancient hybridization event. These results indicate that HGT of *cdtB* occurred about as frequently as *PPO2* duplication events in the sequenced species of Drosophilidae, as CAFE5 predicted 16 *PPO2* duplication events on this tree topology.

**Figure 4.**
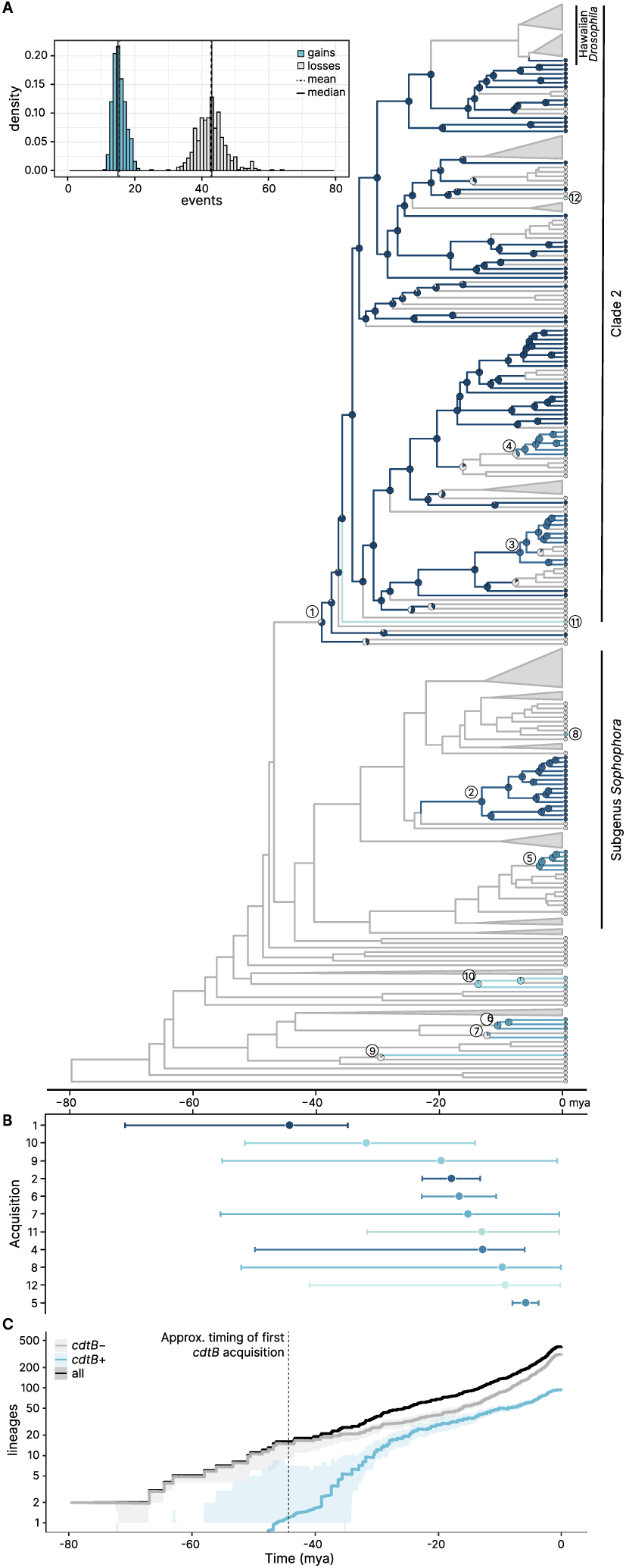
*cdtB* was acquired multiple independent times in the Drosophilidae and is associated with high diversification rates in the early evolution of the genus. (A) Ancestral state reconstructions of *cdtB* acquisitions estimated using total evidence. Circled numbers indicate independent acquisition (or transposition) events according to the scheme in Figure 3B-C, and branches are colored based on dominant ancestral state. Inset summarizes the total number of gain (blue) and loss (grey) events across 500 stochastic character mapping simulations. (B) Mean age for each independent acquisition event as predicted from stochastic character mapping. Error bars indicate 95% confidence intervals. (C) Lineage through time plots showing net diversification of lineages that encode *cdtB* (blue) or do not encode *cdtB* (grey) over time. Ribbons indicate 95% confidence intervals. The y-axis is plotted on a log scale to better visualize changes in diversification rate, but the values themselves are not log-transformed. Detailed views of the ASR phylogeny may be viewed on https://trnpl.github.io/antiparasitoid/.

We then estimated the timing for each predicted independent *cdtB* acquisitions. *cdtB* acquisitions occurred repeatedly throughout the evolution of the Drosophilidae. The oldest acquisition occurred ∼44.3 mya (95% CI: 67.45–34.58 mya, Table S5), before the radiation of subgenus *Drosophila,* and the most recent occurred ∼5.95 mya (95% CI: 8.15–3.84 mya, Table S5) in the *saltans* group (Figure 4B). Several *cdtB* acquisitions occurred within the last 20 million years, including two secondary gain events in the *cardini* group (acquisition 4) and in *D. incompta* (acquisition 12, Figure 4B, Table S5). The five newest *cdtB* acquisitions are all three-exon *ananassae*-type *cdtB* homologs, indicating that this newer copy has a fitness advantage over the subgenus *Drosophila*-type *cdtB*.

### Horizontal transfer of *cdtB* is associated with higher diversification rates in the Drosophilidae

Parasitoid wasps impose strong selective pressure on host populations because they obligately kill their hosts before they reach reproductive maturity. Given the protective effect of gaining *cdtB*, we hypothesized that lineages with *cdtB* were more likely to escape parasitoid infection and would thus experience higher rates of diversification than lineages without *cdtB*. To address this, we first performed sister-clade analysis using six independent *cdtB* acquisition events in Drosophilidae (38). We found an increase in diversification rate associated with *cdtB* acquisition among these pairs of clades (p = 0.017), an effect primarily driven by the subgenus *Drosophila* (Table 1). Phylogenetic generalized least squares (PGLS) regression analyses indicated a trend toward a positive association between *cdtB* presence and net diversification rates under three extinction scenarios (p > 0.20, Table S7, see Materials & Methods). No such relationship was found with *PPO* duplications across five sister-clade comparisons and so we did not conduct PGLS (p = 0.866, Table 1).

**Table 1.** *cdtB* acquisition, but not *PPO* duplication, is associated with higher diversification in the Drosophilidae. Sister-clade analyses were conducted using all pairs. We did subsequent analyses removing one sister pair for *cdtB* to determine the contribution of each group to the effect observed in the full analysis. P-values represent one-sided tests with a null hypothesis µ ≤ µ0. * = p < 0.05, ** = p < 0.01. Further details on methodology and sister-clade selection can be found in Materials & Methods and Table S6.

| Gene | Sister pair removed | Chi-square ($\chi^2$) | Degrees of freedom | P-value | Significance |
| --- | --- | --- | --- | --- | --- |
| ***cdtB*** | None | 4.53 | 1 | 0.014 | * |
|  | Subgenus *Drosophila*+paraphyletic genera/<br>Subgenus *Dorsilopha*+*Microdrosophila* | 0.06 | 1 | 0.401 |  |
|  | *ananassae* subgroup/*setifemur* subgroup | 3.33 | 1 | 0.034 | * |
|  | *saltans* subgroup/*cordata*+*elliptica* subgroups | 5.20 | 1 | 0.011 | * |
|  | *cardini* group/*guarani* group | 8.24 | 1 | 0.002 | ** |
|  | *Leucophenga*/*Amiota*+*Cacoxenus*+*Phortica* | 3.26 | 1 | 0.035 | * |
|  | *Rhinoleucophenga*/*Gitona* | 3.10 | 1 | 0.039 | * |
| ***PPO*** | None | 2.25 | 1 | 0.866 |  |

We next visualized how *cdtB* acquisition was associated with diversification rates over time with lineage through time analyses (see Materials & Methods) (39). *cdtB*-positive (*cdtB*+) lineages experienced higher rates of diversification in the early evolution of the Drosophilidae compared to *cdtB*-negative (*cdtB*-) lineages (Figure 4C). Indeed, the number of *cdtB*+ lineages nearly matched that of *cdtB*-lineages by ∼25 mya, despite the ∼27 million year diversification lead time in *cdtB*- lineages. Following this initial burst in *cdtB*+ lineages, both *cdtB*+ and *cdtB-* lineages diversified at similar rates ∼24–16 mya. Although speculative, the initial fitness advantage conferred by *cdtB* may have been lost to an extent due to resistance co-evolving in parasitoids or other unknown factors that slowed diversification rates. Such resistance to CdtB’s effects was observed in *Leptopilina victoriae*, a specialist parasitoid of *D. ananassae*, which is far less susceptible to its effects than even widely virulent congeners like *L. heterotoma* (21).

Following this interval, *cdtB*- lineages began diversifying at higher rates than *cdtB*+ lineages (Figure 4C). This increase in *cdtB*- diversification rate coincides in part with radiations within the subgenus *Drosophila* that lost *cdtB*. These comprise some of the most diverse species groups in the genus, including the Hawaiian *Drosophila* (620 species), the *repleta* group (173 species), and the *immigrans* group (123 species). These radiations are associated with major biogeographical shifts, including the colonization of the Hawaiian Islands and of new niches like cactus rots in the deserts of the Americas and fungal fruiting bodies, including the toxic, α-amanatin-producing mushrooms that host many species in the *immigrans* radiation (40). Such colonization events in geography and larval breeding niche, or both, may have facilitated a temporary escape from parasitoid pressure. We hypothesize that such releases from parasitoid selection pressure may have coincided with the loss of *cdtB*, either through selection against it arising from a fitness cost (e.g., autoimmunity *sensu* Tarnopol et al. 2025) or a relaxation of purifying selection in the absence of parasitoids (23).

## Discussion

Insects have evolved potent defense mechanisms to resist parasitoids (7, 15, 29). Among the known immune mechanisms, we found that the canonical melanotic encapsulation response to parasitoid challenge is relatively rare in the Drosophilidae and that prophenoloxidase gene duplication events that underlie this response are largely restricted to species in subgenus *Sophophora*. In contrast, we found evidence for at least 11 independent acquisitions of *cdtB* in the Drosophilidae, including before the ancient radiation of the subgenus *Drosophila* ∼44.3 mya and as recently as ∼5.95 mya in *saltans* group species. We estimate that up to 682–2365 drosophilid species encode *cdtB* and up to 1385–2895 species encode *PPO* duplications, though >50% of the latter are accounted for by species in subfamily Steganinae with ancestral *PPO* duplications. Our results suggest that HGT is a surprisingly important source of novelty in insect immune systems against parasitoids, rivaling *PPO* gene duplication.

Although correlational, *cdtB* acquisition was associated with higher diversification rates in the early evolution of the family. Indeed, our findings are consistent with Red Queen-like dynamics wherein *cdtB* acquisition facilitated higher rates of diversification in early-branching lineages of Drosophilidae through resistance to parasitoid wasps, followed by a period of lower rates of diversification as parasitoids co-evolved mechanisms to resist *cdtB*, and finally, a secondary, more recent period of diversification associated with loss of *cdtB* as flies colonized new ecological niches and temporarily may have escaped parasitoid pressure, alleviating the selective constraint to retain *cdtB* in the genome. Although speculative, this “escape and radiate” scenario (*sensu* Ehrlich and Raven 1964) aligns with observations from herbivorous insects that switch hosts or become specialized to escape generalist enemies (41, 42). These results highlight the potential for humoral effectors as robust anti-parasitoid molecules that can facilitate rapid adaptation to parasitoid pressure, potentially bypassing more costly developmental programs underlying the evolution of specialized hemocytes.

There are several important limitations to our study. One arises from the lack of empirical data for anti-parasitoid immunity across the Drosophilidae. The anti-parasitoid immune responses are unknown for most species, particularly outside of the subgenus *Sophophora*, which comprise ∼80% of species with phenotyped anti-parasitoid responses. Further studies are needed to determine whether new *PPO* duplications (e.g. *PPO4*) enable convergent evolution of melanotic encapsulation phenotypes and whether *cdtB* truly functions in anti-parasitoid immunity in independent acquisitions outside of the *ananassae* subgroup. Moreover, anti-parasitoid immune mechanisms independent of these genes might also be widespread in the Drosophilidae, such as new encapsulating cell types (43–47), symbiont-mediated immunity (48, 49), behavioral and physiological adaptation to chemical environments (50–52), other horizontally transferred effectors (53), or expansions of clotting factors like ficolins and fibrinogens that are reported here.

These limitations notwithstanding, our study highlights that while model organisms like *D. melanogaster* provide a powerful platform for the discovery of fundamental biological processes, each model organism has its own unique phylogenetic history. Without a broader evolutionary context, derived traits, such as the PPO-based melanotic encapsulation anti-parasitoid defenses of *D. melanogaster*, can instead be interpreted as ancestral (54, 55). Detailed study of non-canonical yet widespread humoral immune effectors against blood and hemolymph-feeding parasitic animals may unveil general principles that underlie anti-macroparasite immune responses in both invertebrates and vertebrates.

## Materials and Methods

### Defense mechanism assignments

To determine the prevalence of anti-parasitoid defense mechanisms described in Drosophilidae, we conducted a literature review for laboratory studies of wasp challenge in drosophilid hosts. We used articles that reviewed anti-parasitoid immune mechanisms in drosophilid species (6, 16, 56) to find appropriate primary articles, then expanded the search to include all works of particular scientists who extensively studied anti-parasitoid immunity in diverse drosophilids (i.e., AJ Nappi, Y Carton, FA Streams, T Schlenke). Subsequent Google Scholar searches using search terms including “Drosophila parasitoid immunity” were then used to identify additional articles that might have been missed. Data were only included for species where wasp infection and subsequent immune reactions were observed in the laboratory, excluding studies that performed laboratory infections and did not investigate immune responses or that characterized putative encapsulating hemocytes in the absence of wasp infection. Species in which more than one defense mechanism was observed were assigned to multiple categories. Defense mechanisms and the studies from which they were derived can be found in Table S8 and Supplementary File S1.

### Genome assembly

All drosophilid species with publicly available genome assemblies on NCBI as of April 2026 were included in this study. Additionally, TBLASTN searches using drosophilid CdtB as queries (see Manual curation and annotation of *PPO* and *cdtB* genes) were conducted on all drosophilid species that did not have genome assemblies but had public NCBI Short Read Archive (SRA) datasets to increase taxonomic sampling in our study. Species that had hits against *cdtB* in their SRA dataset were subsequently assembled for further analysis, as were, if available, those of closely related species where *cdtB* wasn’t initially detected. We also included previously unpublished genomes for *D. trisetosa* and *D. multispina* to fill in taxonomic gaps in subgenus *Drosophila*. A full list of drosophilid genomic resources used in this study can be found in Supplementary File S1.

For species with new, high-accuracy (Oxford Nanopore R10.4.1) long-read sequencing data, genomes were assembled using DNA extraction and sequencing protocols developed by Kim et al. 2024 (57). Briefly, genomic DNA was extracted from single flies by phenol-chloroform extraction and a sequencing library was prepared following the standard ONT LSK114 ligation sequencing kit protocol, scaling volumes by 0.5x for cost efficiency. A complete protocol is provided at: https://dx.doi.org/10.17504/protocols.io.ewov1q967gr2/v1. Sequencing was performed on an ONT P2 Solo and basecalling was performed with Dorado v0.9.6 using the dna_r10.4.1_e8.2_400bps_sup@v5.2.0 model. Genomes were assembled with hifiasm 0.25.0 (58) and the primary assembly was provided to downstream steps. Alternatively, if unassembled short reads were the only available source of data, SPAdes v4.2.0 (59)with default settings was used to generate the assembly. Contaminant sequences were flagged and removed with NCBI Foreign Contamination Screen v0.5.5 (60).

### Species tree construction

A species tree was estimated generally following the phylogenomic workflow used in Suvorov et al. 2022 and Kim et al. 2024 (57, 61). BUSCO v5.8.0 (62) with the diptera_odb10 database was used to annotate conserved single-copy orthologs in all genome assemblies. Coding sequences were extracted for the subset of 1,000 most present orthologs, aligned with MAFFT v7.490 (63) with default settings, and individual gene trees were estimated with IQTREE3 v3.0.1 (64) with the GTR+I+R5 substitution model. Then, a consensus species tree was estimated with ASTRAL v5.15.5 (65). Codons containing four-fold degenerate positions were identified with msa_view from PHAST v1.4 (66), using the *D. melanogaster* annotations as the reference. These sites were extracted, and fixing the previously inferred topology, the branch lengths of the species tree were rescaled by the 4-fold site substitution rate. Finally, the species tree was scaled to chronological time. Labels for the time-scaled phylogeny inferred by Suvorov et al. 2022 (61) were harmonized with this tree. Secondary calibration points were identified using the congruify.phylo method from geiger 2.0.11 (67), using the treePL v1.0 (68) penalized likelihood framework to scale to chronological time.

### Manual curation and annotation of *PPO* and *cdtB* genes

*PPO* and *cdtB* genes were identified in drosophilid genomes using TBLASTN searches (69) with *Drosophila melanogaster* PPO1, PPO2, and PPO3 amino acid sequences (*PPO*), and *D. ananassae*, *D. primaeva*, *D. biarmipes*, *Scaptomyza flava*, and *Zaprionus bogoriensis* CdtB amino acid sequences as queries (*cdtB*, see Supplementary File S1 for accessions). Hits were mapped to relevant contigs, and contigs were pulled for manual inspection. Where available, we used the drosophilid annotation dataset curated by Dhakad et al. 2026 (70). For new species that were not included in this dataset, we used either LiftOn v 1.0.2 (71) to port annotations from closely related species to serve as baseline annotations or Augustus (72) as implemented in Geneious Prime 2026.1 using the “fly” model to predict annotations. These annotations were manually inspected and, if necessary, edited. Gene neighborhoods were determined through BLASTP searches of neighboring genes. Further details on annotation can be found in *SI Methods*.

### Gene tree construction

To generate gene trees for *PPO* homologs, nucleotide sequences for the coding regions of all intact drosophilid *PPO* genes annotated in this study as well as select outgroup insect and crustacean *PPO* genes were aligned and trimmed using the MACSE OMM pipeline (73). Both nucleotide and translated amino acid alignments are produced. These alignments were run in IQTree v 3.1.2 (64), where tree topologies were estimated using the best fit model determined by ModelFinder Plus (74) with 1000 ultrafast bootstraps replicates (75) and 1000 SH-aLRT replicates (76) to assess branch support. The tree was rooted with crustacean *PPO* homologs as outgroups. The same pipeline was used to generate gene trees for drosophilid *PPO1* and *PPO2/3/4* homologs for molecular evolution analyses within these orthology groups.

*cdtB* homologs are not orthologous between free-living bacteria and insects, which inhibited nucleotide-based phylogenetic inference. Instead, an amino acid alignment comprising the drosophilid CdtB homologs identified in this study and those annotated in public databases was used to infer relationships between CdtB homologs. Additional insect and bacterial CdtB homologs were included through BLASTP searches of the NCBI nr database using *D. ananassae* and *S. flava* CdtB as queries (see Supplementary File S1 for accessions). The list of hits was reduced according to criteria described in *SI Methods*, aligned using MAFFT v 7.526 (63), and trimmed from ends manually and then with the smartgap algorithm on the Clipkit webserver (77). An amino acid tree was constructed using IQTree v 3.1.2 (64) using the same settings as described above. The tree was then rooted with the clade comprising free-living bacterial CdtB homologs as the outgroup.

### Gene family evolution analyses

To model the rate of *PPO* gain and loss in the Drosophilidae, we ran CAFE5 with the ultrametric species tree we constructed for this study (78). Our PPO gene copy number counts included the PPOs we had evidence for (i.e., poor assemblies and partial PPOs) but excluded pseudogenizations and likely misassemblies. Because CAFE5 requires the root node to have at least one copy of each paralog, we modeled *PPO2, PPO3,* and *PPO4* as one gene family and *PPO1* as another. We did not include *cdtB* in our CAFE5 analyses because we predicted *cdtB* was not present at the root node of our phylogeny (see Ancestral state reconstruction for HGT events). CAFE5 analyses were conducted using the gamma model with two rate categories (k = 2), which were a better fit for the data than the base model (Likelihood-ratio test, p=0.014).

### Protein evolution analyses

For tests of positive selection, we generated separate subtrees comprising *PPO1* orthologs and *PPO2/3/4* orthologs from Drosophilidae and *Liriomyza trifolii* PPOs as outgroups only using the MACSE OMM pipeline as described above. These two trees were used for further analysis with the HyPhy package for comparative sequence analysis (79). To identify if there was evidence of episodic positive selection in each of the *PPO* clades, we ran BUSTED with default settings (80). For both the *PPO2/3/4* tree and *PPO1* tree, we set foreground (test) branches to the combined *Drosophila* and descendent genera (e.g. *Scaptomyza*, *Lordiphosa*, etc.) clade (including the branch leading into this clade), which decreases the chance of false positives by testing only a target subset of flies instead of the entire tree. We tested each of the three major *PPO2* duplication events (ancestral duplication in basal drosophilids, *PPO3*, and *PPO4*) including the branch leading into each clade as the foreground against the rest of the *PPO2/3/4* tree to test if these duplication events were associated with positive, diversifying selection.

Since we found evidence of diversifying positive selection for all three *PPO2* duplication clades, we subsequently ran a branch-site test using aBSREL (81) with each set as foreground again to identify specific branches that contributed to signals of positive selection. To show that the evidence of positive, diversifying selection was specific to the evolutionary histories of duplicated *PPO*s, rather than *PPO2*, we reran aBSREL with the sister *PPO2* clade set as the foreground (i.e. the clade consisting of ‘oriental’ group species’ *PPO2* orthologs for *PPO3* and the clade consisting of *saltans* and *willistoni* species groups’ *PPO2* orthologs for *PPO4*) and the rest of the tree as background. We could not make the sister comparison for the ancestral *PPO2* duplication in basal drosophilids because the sister clade was not monophyletic. We also did not further test the *PPO1* tree because we had no specific hypotheses.

We did not conduct tests for diversifying selection on *cdtB* homologs because they are not orthologous, and as such we would expect elevated dN/dS due to nucleotide diversity that stems from independent origins of each copy, rather than from true signals of positive selection.

### Pancrustacea *PPO* copy number inference

To quantify the extent of *PPO* copy number duplications across insects and crustaceans, we screened all RefSeq annotated genomes available on NCBI for all insects and crustaceans that had genomic resources available. We searched the annotation dataset for “phenoloxidase” and counted only entries annotated as “phenoloxidase,” “phenoloxidase-like,” or “phenoloxidase subunit-like.” We excluded species where annotations did not include functional annotations or where the phenoloxidase search returned no results since we were unable to perform manual quality control analyses to confirm true absence of *PPO* genes in these taxa. To find the average *PPO* copy number in Pancrustacea, we averaged the mean copy number in each insect order and crustaceans to control for variance in genome resource availability across orders. A full list of species included in the analysis can be found in Supplementary File S1.

### Ancestral state reconstruction for HGT events

Ancestral state reconstruction was conducted using the fitMk function in phytools v 2.4.4 using custom transition matrices to fit a “total evidence” model where synteny and gene phylogeny information were used to assign distinct states to putative independent acquisition events (82). Acquisitions were considered independent if both of the following criteria were met: 1) *cdtB* is encoded in a different syntenic locus and 2) the *cdtB* homolog is not most closely related to those found in sister species on the species tree. An acquisition was classified as a transposition event if criterion 1 was met but criterion 2 was not. For taxa that encoded *cdtB* on short contigs lacking synteny information, we assigned it as an independent acquisition if the placement of its CdtB homolog in the amino acid phylogeny was discordant with the placement of the taxon in the species phylogeny. Using these criteria, we identified 12 states comprising 11 independent acquisition events and one transposition event between two loci in *Zaprionus*.

For each model, we made the following assumptions: 1) gain events were rarer than loss events; 2) with the exception of one probable transposition event in *Zaprionus*, transitions between syntenic loci did not occur; and 3) *cdtB* was not present in the last common ancestor between Drosophilidae and Agromyzidae. To address assumptions 1 and 2, our transition matrix modeled gain and loss events under distinct rates for each state, and prohibited transitions were given a rate of 0. To address assumption 3, we set the root prior to a 100% probability of *cdtB* absence. The models from fitMk were then passed to the ancr function in phytools to calculate ancestral states at each node. For clarity, only nodes where P(no *cdtB*) ≤ 0.90 were mapped.

The fitMk models were also passed to the make.simmap function in phytools for stochastic character mapping to estimate the number of gain and loss events across the tree and the dominant state along each branch of the tree. 500 simulations were run for each model. The countSimmap function summarized the number of gains and losses for each state across the species tree. Gains and losses inferred for each state transition using countSimmap were combined for each simulation to find the distributions of total gains and losses predicted by this model.

### Dating *PPO* duplication and *cdtB* acquisition events

*PPO* duplication events were dated using ancestral states calculated through CAFE5 analysis. Since CAFE5 does not provide a simulation output, we estimated duplication time based on the age of the node inclusive of the last common ancestor that shared the duplicated state.

To infer the timing of each independent *cdtB* acquisition event, we recorded the earliest age at which each acquisition state was observed from each simulated stochastic character map. 95% confidence intervals were calculated by determining the values comprising the upper and lower 2.5% quantiles of acquisition ages across all simulations.

### Diversification analyses

#### Sister-clade analyses

For sister-clade analyses comparing diversification rates between clades that encode *cdtB* and those that do not, we used sister clades representing independent acquisitions of *cdtB*. We excluded any independent acquisition from “one-off” species where ASR did not predict presence of that *cdtB* acquisition at the node with its sister taxon (i.e. *D. biarmipes, D. incompta,* and *Dettopsomyia nigrovittata*). We excluded *Colocasiomyia* and its sister clade from this analysis due to discordances in our tree topology and those reported in Dias et al. 2025 (83), which creates uncertainty in both stem age and species richness in each sister clade. We used the same general principles to find sister pairs for *PPO* duplication events. This resulted in a test comprising six sister pairs for *cdtB* and five sister pairs for *PPO* duplications (Table S6).

Species richness for each clade was inferred from the number of described species in NCBI Taxonomy for each taxonomic group. Stem ages for clades were taken from Suvorov et al. 2022 (61) if the clades were within Drosophilinae and from Dias et al. 2025 (83) if they were outside Drosophilinae . Species richness and stem ages for each pair are reported in Table S6. The sister-clade test was conducted using the richness.yule.test function in ape v 5.8.1 (84), and the p-value was corrected for a one-sided test for the null hypothesis µ ≤ µ_0_. For *cdtB*, we tested for the contribution of any one sister pair by conducting additional tests with one pair removed.

#### Diversification rate phylogenetic generalized least squares regression

To follow up on the relationship between *cdtB* presence and net diversification rate, we performed phylogenetic generalized least squares (PGLS) regression using the methodology described in Wiens et al. 2015 (85) and Guo et al. 2023 (86). Briefly, net diversification rates were calculated under three extinction scenarios (0% extinction, 50% extinction, and 90% extinction) using the <u>bd.ms</u> function in the R package geiger v 2.0.11 (87). We then generated a reduced tree that comprised one taxon per clade included in the sister pair analysis and tested for a relationship between *cdtB* presence in the clade (independent variable) and net diversification rate (dependent variable) using PGLS in the R package caper v 1.0.3 (https://github.com/davidorme/caper). Lambda parameters were inferred by the model, while kappa and delta were fixed at 1. Input data for the PGLS analysis can be found in Table S9.

#### Lineage through time (LTT) analysis

LTT plots are useful for visualizing shifts in diversification through time, where shifts in slope indicate increases or decreases in diversification rate (39). To generate LTT plots segregated by character state of branches, we used the stochastic character mapping data to track the average number of branches (lineages) that were *cdtB*-positive (*cdtB*+) or *cdtB-*negative (*cdtB*-) over time, following the tutorial here: https://blog.phytools.org/2022/08/lineage-through-time-plots-for.html. Branch states were computed from 500 simulated trees with singleton nodes computed using the map.to.singleton function in phytools v 2.4.4 such that each edge only had one state. 95% confidence intervals were calculated by finding the upper and lower 2.5% quantile lineage values at a given time t where the number of lineages ≥ 1.

### Data and Materials Availability

A browsable webserver for all phylogeny-wide analyses can be accessed at https://trnpl.github.io/antiparasitoid/. Code for phylogenetic analyses and figure generation and full resolution versions of main text figures can be found at the corresponding github repository: https://github.com/trnpl/antiparasitoid/tree/main/code. All other raw data associated with the manuscript (phylogenies, alignment files, and annotated contigs) can be accessed from the Dryad repository for this project: https://doi.org/10.5061/dryad.s7h44j1qm

## Acknowledgments

This study was supported by NIH award #R35GM119816 to N.K.W. We thank Artyom Kopp for providing samples for genome assemblies for *Drosophila trisetosa* and *D. multispina.* We also thank Diler Haji and Jeffrey Groh for helpful comments and advice on analyses performed in this manuscript.

## Author Contributions

R.L.T. and N.K.W. conceptualized the study. B.Y.K. provided genome assemblies and the ultrametric species tree comprising all genomes surveyed in this study. R.L.T. and R.L.W. curated manual annotations and performed all subsequent analyses. R.L.T. visualized the data and built the phylogeny browser. R.L.T. and N.K.W. supervised the project and wrote the first draft of the manuscript. All authors contributed to revisions and editing of the manuscript.

## Supporting information for

### SI Methods

#### Manual curation and annotation of *PPO* and *cdtB* genes

Existing annotations for *PPO* genes on GenBank frequently misassign *PPO* paralogs (i.e., a gene can be annotated as “phenoloxidase 2-like” when it is orthologous to *PPO1*), likely due to the use of automated annotation software. We initially assigned *PPO1, PPO2,* and *PPO3* labels based on conserved synteny throughout Drosophilidae, validated through BLASTP searches of neighboring genes. For *PPO* orthologs that were in new genomic locations or on short contigs that lacked neighboring genes, PPO identity was assigned based on gene phylogenies (see Gene tree construction, Materials & Methods). Due to their deep conservation across Drosophilidae, taxa that were missing either *PPO1* or *PPO2* from the initial BLAST search were subject to a second TBLASTN search to double-check for true loss. Additionally, *PPO* hits were reciprocally BLASTed back to ensure the hit ID was a *PPO* and not *larval serum protein* paralogs, which shared enough sequence similarity to be frequently recovered by BLAST. Manual inspection referenced *PPO* orthologs from RefSeq genomes with RNA-seq supported genome annotations. Manual editing involved predicting introns by applying the GT-AG rule, while allowing for alternative-motif exceptions such as GC-AG (1), and aligning against the representative species’ annotated sequences using MAFFT (2).

Since *cdtB* is not included in the fly BUSCO set, there frequently were not annotations at the *cdtB* locus. In the cases where *cdtB* was predicted, the predicted annotation was used as a starting point to annotate the locus. Through these annotations we serendipitously discovered new intron-bearing *cdtB* structures in some lineages, including *Scaptomyza flava*, which we previously thought did not bear introns (3, 4). When possible, we validated predicted introns against existing transcriptomic datasets in the NCBI SRA, using BLASTN with the predicted *cdtB* sequence and the sequence of some neighboring genes as a query to pull localized reads from the dataset. Reads were then mapped back to the locus in Geneious Prime v. 2026.1 and surveyed for gaps spanning the predicted intron. For taxa that lacked transcriptomic data and/or annotated *cdtB,* we took a parsimony-based approach to annotate the *cdtB* locus, where annotated *cdtB* homologs from closely related species were aligned as a reference using CLUSTALomega v 1.2.3 (5) as implemented in Geneious v 2026.1. The structures for new *ananassae*-like *cdtB* copies were resolved using initial predictions produced by the “fly” or “honeybee” models in Augustus as implemented in Geneious Prime. These predictions were then manually inspected and edited to retain, if possible, the conserved splice junctions previously observed in *D. ananassae* and *D. biarmipes*. These splice junctions were validated using the Berkeley Drosophila Genome Project Splice Site Predictor (https://www.fruitfly.org/seq_tools/splice.html). The BDGP splice site predictor was also used to predict likely splice junctions for divergent three-exon copies of *cdtB*, such as those found in the *saltans* group.

To distinguish putative pseudogenization events from assembly errors, *PPO* and *cdtB* sequences that had predicted reading frame errors by mapping raw genomic reads from available NCBI SRA datasets to correct possible assembly errors. We manually corrected the genomic sequence in cases where assembly errors were observed. In some cases, genome assembly quality prohibited the identification of full-length genes. Strings of ambiguous base calls (Ns) that still went unresolved based on SRA data were marked as “poor assemblies.” Some assemblies had short contigs containing partial gene sequences, or in the case of *cdtB-*bearing species identified through a small number of transcriptome SRA hits, did not include a contig with *cdtB*. We coded these species as having “evidence for” these genes, though their exact sequences could not be deduced. Copy number was often impossible to deduce for these orthologs, so we default assigned one copy of each for gene count analyses. These orthologs, along with pseudogenized copies, were excluded from subsequent molecular evolution analyses.

#### Gene tree construction

##### CdtB sequence representation

BLASTP searches for CdtB in the nr database yielded >1000 hits. To pare down the number of representative hits included, redundant BLAST hits were first removed using cdhit v 4.8.1 (6) at a 97% identity threshold. To improve the alignment, bacterial BLAST hits were further filtered using the following criteria:

1. E-values < 1e-5
2. Total length > 150 and < 400 amino acids
3. Sequences contained the canonical DNAse-I SDH motif

All insect CdtB homologs were retained in the final dataset. The final dataset contained 353 sequences, which were aligned using MAFFT v7.526 (2). The initial alignment was then manually trimmed at poorly aligned N and C termini until a site with ∼70% occupancy was reached and realigned using MAFFT twice. The manually trimmed alignment was further trimmed using the smartgap algorithm on the Clipkit webserver (7). The final alignment comprised 607 sites.

**Figure S1.**
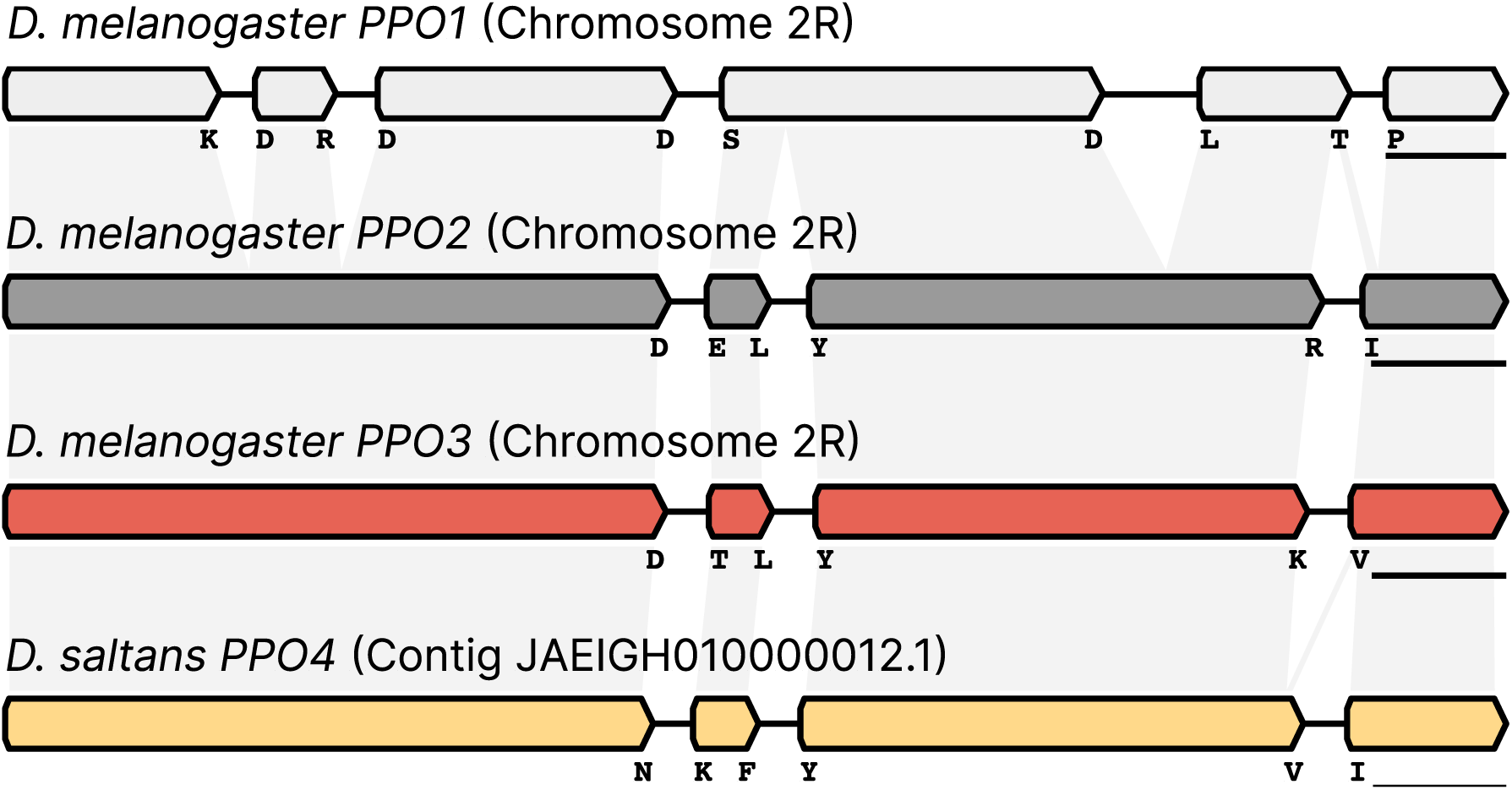
*PPO* gene structures, related to Figure 2. *PPO3* and *PPO4* share a gene structure with *PPO2*. Letters underneath the cartoons indicate residues flanking splice junctions, and grey boxes indicate homology between paralogs. Scale bar = 200bp.

**Figure S2.**
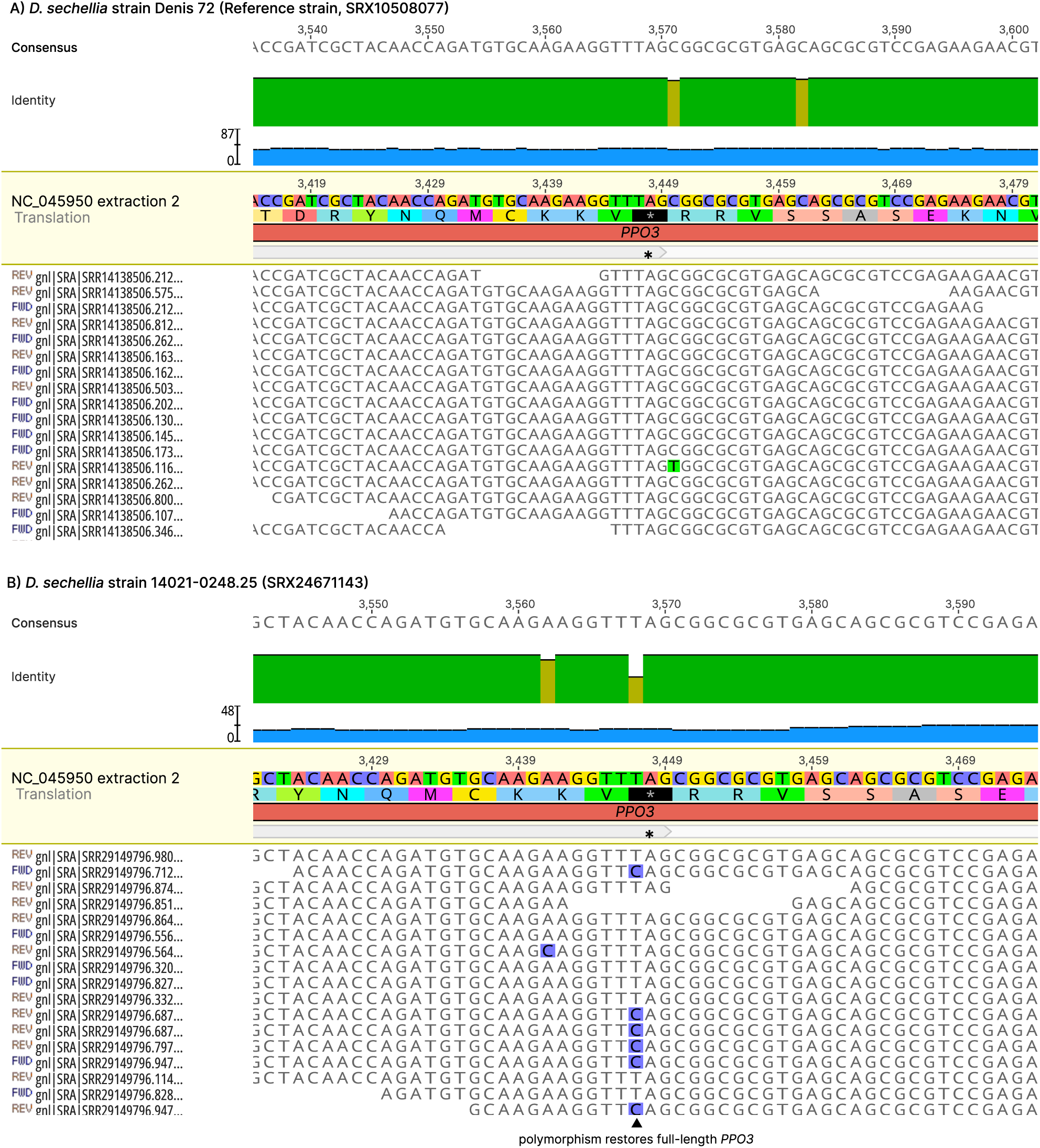
*D. sechellia* is polymorphic for pseudogenized *PPO3,* related to Figure 2. Genomic short reads mapped to *D. sechellia* Denis72 reference strain (A) and *D. sechellia* strain 14021-0428.25 (B) demonstrate segregating polymorphism for pseudogenized and functional *PPO3* alleles.

**Figure S3.**
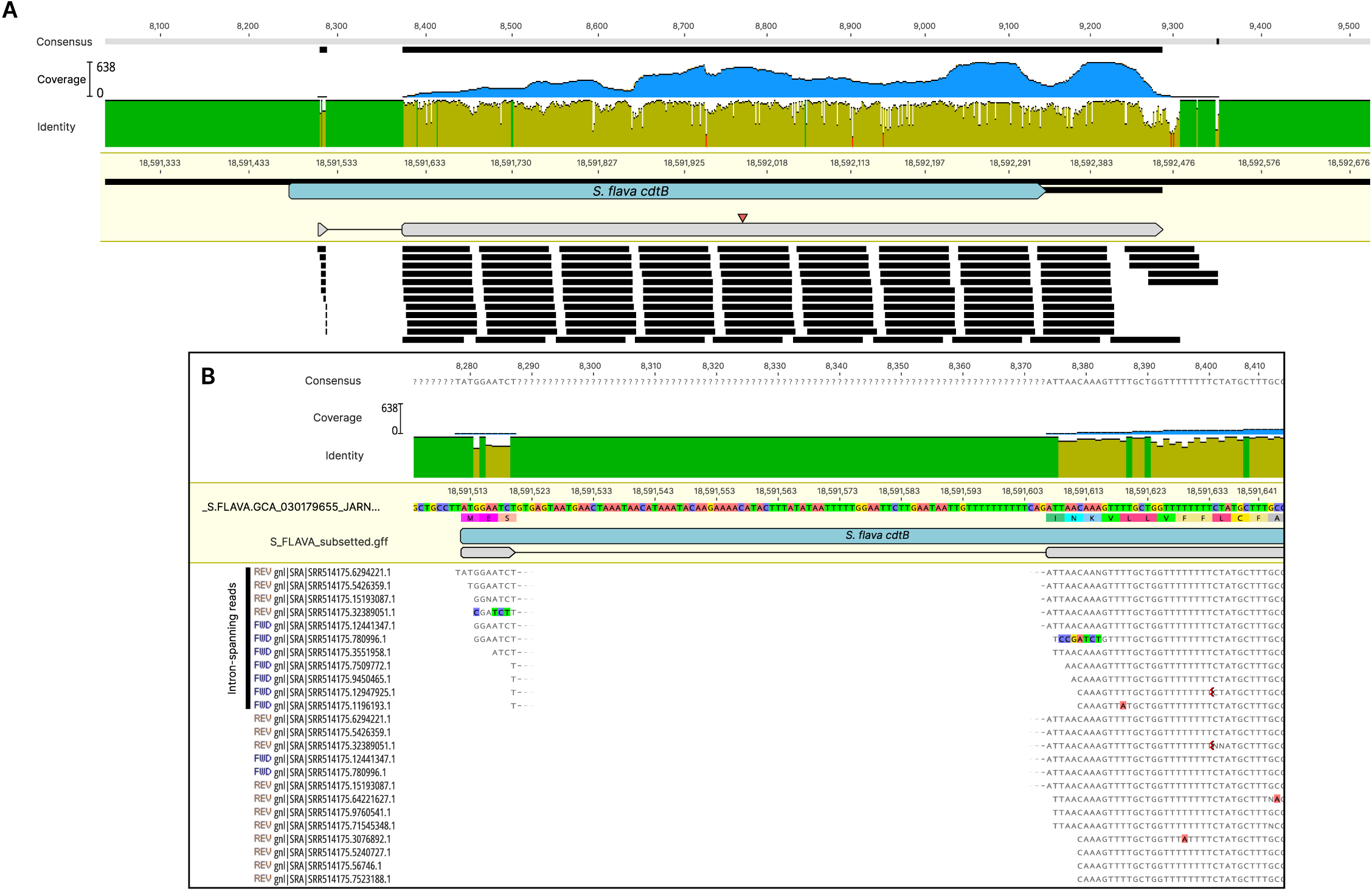
RNAseq validates cryptic introns in *Scaptomyza flava cdtB,* related to Figure 3. (A) RNAseq reads mapping to *S. flava cdtB* extend the previously predicted open reading frame. The red triangle denotes the start codon of the previously predicted ORF. (B) Inset of reads supporting an intron in the *S. flava cdtB* transcript.

**Figure S4.**
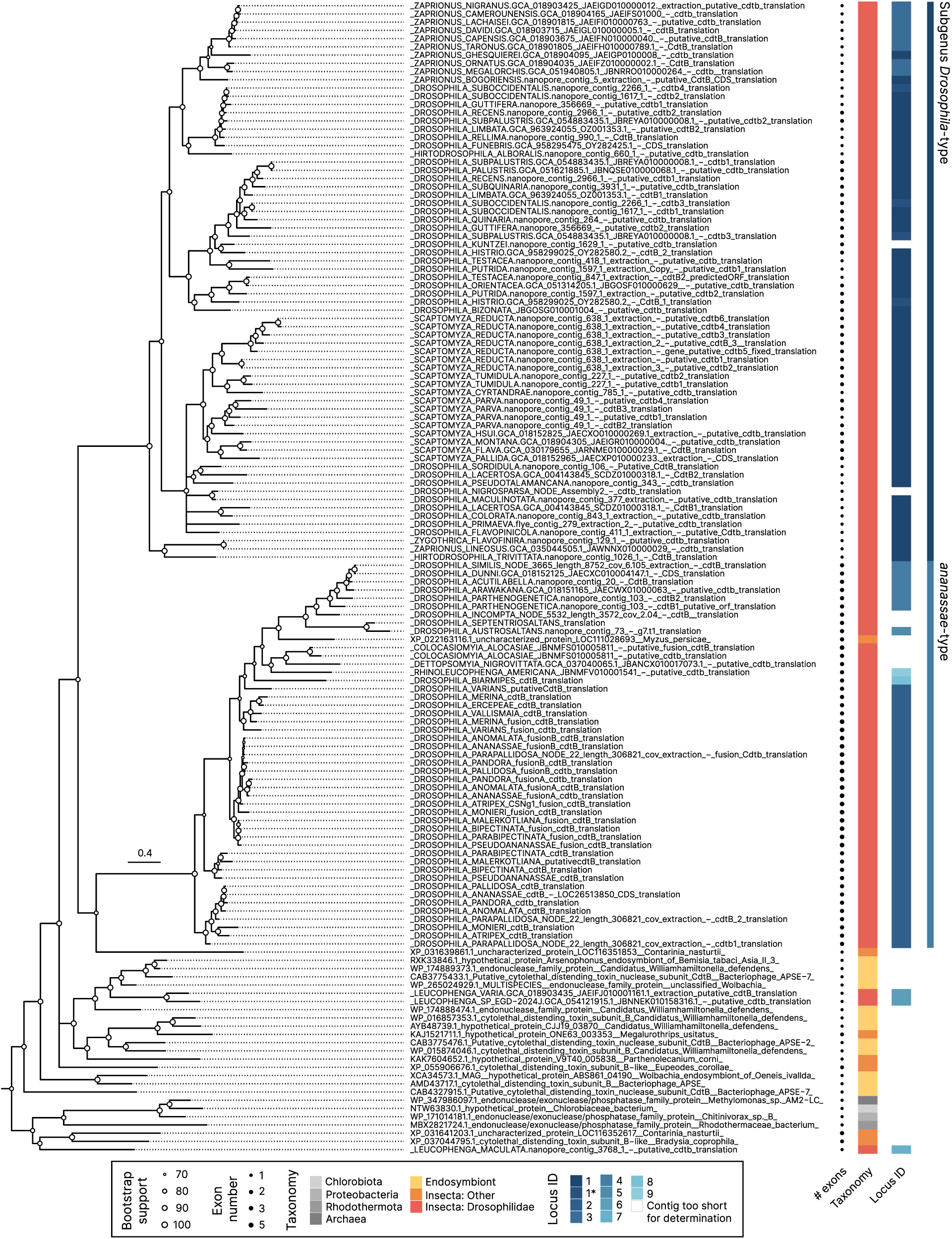
Drosophilid CdtBs form two major clades, related to Figure 3. Insect and endosymbiont CdtB homologs are closely related to insect copies of CdtB. Nodes with < 50% bootstrap support were collapsed into polytomies. Scale bar = substitutions/site.

**Figure S5.**
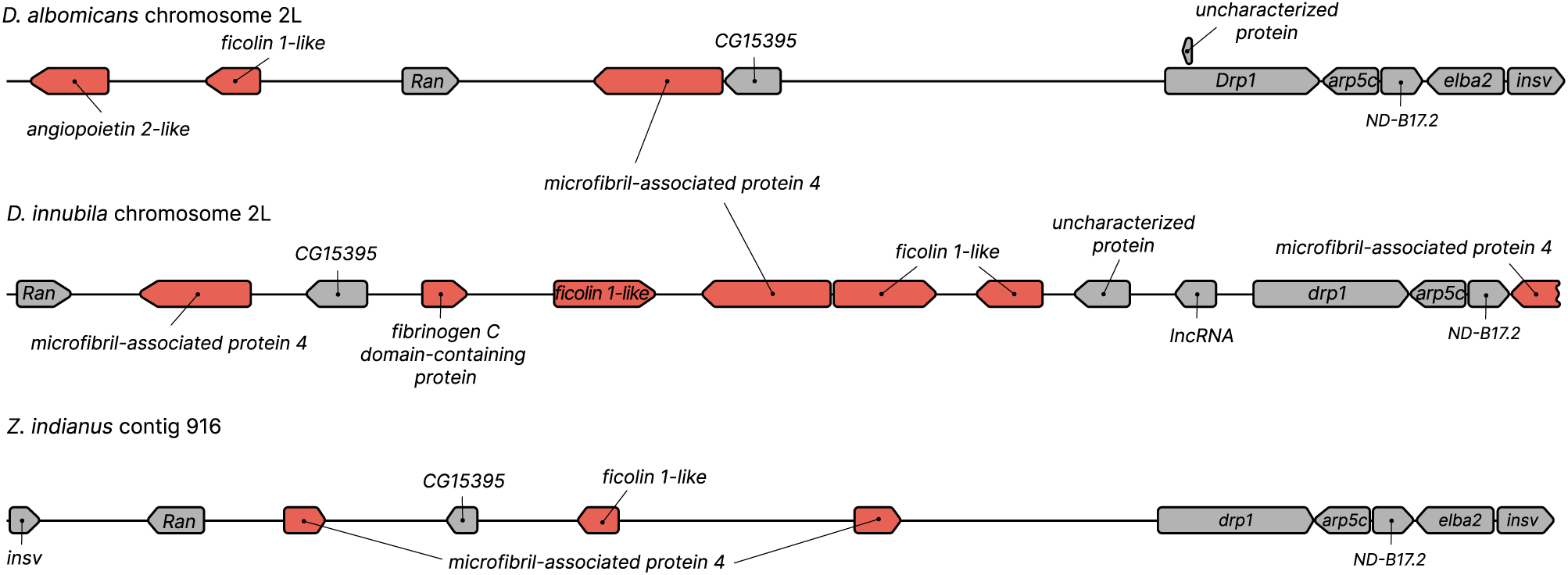
Expansion of fibrinogen-like genes in subgenus *Drosophila* species, related to Figure 3. Species that do not encode *cdtB* retain multiple copies of fibrinogen-like genes in the locus where *cdtB* is frequently found. Cartoon gene schematics are not to scale, but reflect relative gene richness between the conserved *drp1* and *Ran* loci in each species.

**Table S1.** CAFE5 rate classes and results for *PPO1* and *PPO2/3/4* gene family extractions, related to Figure 2C.

| Gene family | Gamma Cat Mean | Posterior Probability | Family-wide P-value<br>(significant expansion/contraction) |
| --- | --- | --- | --- |
| *PPO1* | 0.544602 | **0.999306\*** | 0.999 |
| *PPO1* | 1.4554 | 0.000694385 |  |
| *PPO2/3/4* | 0.544602 | 9.80E-06 | 0.997 |
| *PPO2/3/4* | 1.4554 | **0.99999\*** |  |

**Table S2.** HyPhy results for positive selection in *PPO1* and *PPO2/3/4* orthology groups using BUSTED and aBSREL. Bolded values indicate mean ω > 1.

| Test clades | BUSTED results |  |  |  |  |  | aBSREL results |  |  |  |  |  |  |  |
| --- | --- | --- | --- | --- | --- | --- | --- | --- | --- | --- | --- | --- | --- | --- |
| | p-value | Log(L) | AIC-c | # parameters | Mean $\omega$ (test) | Mean $\omega$ (back-ground) | p-value | Significant branches | Mean $\omega$ | Branch p-value | p-value (Sister *PPO2*) | Significant branches | Mean $\omega$ | Branch p-value |
| PPO1, all foreground | 0 | -153971 | 309484 | 769 | 0.1049 | NA |  |  |  |  |  |  |  |  |
| PPO1–*Drosophila* | 0 | -153936 | 309424 | 774 | 0.1016 | 0.1421 |  |  |  |  |  |  |  |  |
| PPO2, all foreground | 0 | -187870 | 377646 | 950 | 0.07721 | NA |  |  |  |  |  |  |  |  |
| PPO2–*Drosophila* | 0 | -187849 | 377613 | 955 | 0.07095 | 0.09204 |  |  |  |  |  |  |  |  |
| PPO2—Basal Drosophilidae duplications | 5.45E-09 | -187857 | 377630 | 955 | 0.1146 | 0.07268 | p≤0.05 | Node 869 (stem to all *PPO2* in this clade) | **2.814** | 0 |  | Not tested (sister *PPO2*s not monophyletic) |  |  |
|  |  |  |  |  |  |  |  | Node 905 (stem leading to non-*Drosophilini* *Drosophilinae* *PPO2*) | **1.552** | 0.002 |  | Not tested (sister *PPO2*s not monophyletic) |  |  |
|  |  |  |  |  |  |  |  | Node 870 (stem leading to *Steganinae* *PPO2*) | **72.09** | 0.029 |  | Not tested (sister *PPO2*s not monophyletic) |  |  |
| PPO3 | 6.94E-10 | -187108 | 376132 | 955 | 0.2894 | 0.06367 | p≤0.05 | Node 81 (stem to all *PPO3*s) | **1.933** | 0.001 | p≤0.05 | Node 178 (stem for *D. erecta* and *D. oreana* *PPO2*) | 0.4038 | 0.034 |
| PPO4 | 7.15E-5 | -187602 | 377120 | 955 | 0.3636 | 0.07141 | p≤0.05 | Node 16 (stem of all *PPO4*s) | **1.427** | 0.01 | p≤0.05 | None | NA | NA |
|  |  |  |  |  |  |  |  | Node 29 (stem to *sturtevantii* subgroup *PPO4*) | 0.7671 | 0.018 |  |  |  |  |

**Table S3.** Average *PPO* copy number in drosophilid clades, other Diptera, and other insects and crustaceans, related to Figure 2D. P-values reflect differences from the pancrustacean average via two-sample t-tests with Bonferroni correction. For species-specific counts, see Supplementary File S1.

| **Taxon** | **# species** | **Average *PPO* count (± S.D.)** | **Adjusted p-value** |
| --- | --- | --- | --- |
| Drosophila Clade 1 | 139 | 2.417266 ± 0.7009236 | 1.1418E-12 |
| Drosophila Clade 2 | 230 | 2.052174 ± 0.2416637 | 0 |
| Basal Drosophilidae | 37 | 2.864865 ± 1.158595 | 0.017166 |
| Non-drosophilid Diptera | 72 | 5.916667 ± 3.0569708 | 0.00059748 |
| Other insects | 246 | 2.617886 ± 2.366502 | 2.6646E-15 |
| Crustaceans | 24 | 1.791667 ± 1.1025333 | 0.00028782 |
| Pancrustacea average<br>(normalized across orders) | 748 | 4.410497 ± 3.128569 | |

**Table S4.** Summary statistics for 500 simulations of *cdtB* acquisition ancestral state reconstruction, related to Figure 4A.

| Model | Mean gains<br>(± S.D.) | Median<br>gains | Minimum<br>gains | Maximum<br>gains | Mean losses<br>(± S.D.) | Median<br>losses | Minimum<br>losses | Maximum<br>losses |
| --- | --- | --- | --- | --- | --- | --- | --- | --- |
| Total<br>evidence | 15.428 ± 1.929 | 15 | 11 | 25 | 42.81 ± 4.393 | 43 | 30 | 64 |

**Table S5.** *cdtB* acquisition timing based on ancestral state reconstruction simulations, related to Figure 4B. Timing of acquisition 3 was not calculated since this was a likely transposition event, rather than an independent acquisition event.

| Acquisition event | Mean age (million years) | 95% CI upper bound age (million years) | 95% CI lower bound age (million years) | Maximum age (million years) |
| --- | --- | --- | --- | --- |
| 1 | 44.32019215 | 67.45227904 | 34.58435167 | 78.67811071 |
| 2 | 18.10982437 | 22.69980796 | 13.31907225 | 22.94745998 |
| 4 | 12.18541602 | 51.94251957 | 6.116803609 | 78.09112095 |
| 5 | 5.946497291 | 8.153835226 | 3.835012025 | 8.220590852 |
| 6 | 16.62573112 | 22.66984267 | 10.79757423 | 23.16270874 |
| 7 | 15.33017666 | 54.08856812 | 0.2889832654 | 78.2809012 |
| 8 | 11.40844165 | 61.14693282 | 0.2094189948 | 78.2957867 |
| 9 | 20.03576727 | 49.81874042 | 1.151788632 | 71.6937776 |
| 10 | 31.46818016 | 51.62587416 | 14.09512529 | 66.40192619 |
| 11 | 13.67959374 | 32.18388543 | 0.9036045112 | 35.27825447 |
| 12 | 9.427411724 | 38.90209089 | 0.1983422278 | 78.14199072 |

**Table S6.** Sister-clade pair stem age and species richness, related to Table 1. Species richness was estimated from NCBI Taxonomy counts for the respective taxonomic divisions.

| Sister pair | Crown clade with derived state | Gene | Species richness of derived clade (n) | Crown clade with ancestral state | Species richness of ancestral clade (n) | Divergence time (my) | Divergence time reference |
| --- | --- | --- | --- | --- | --- | --- | --- |
| Pair 1 | Subgenus *Drosophila* + paraphyletic genera | *cdtB* | 3079 | Subgenus *Dorsilopha* + *Microdrosophila* | 107 | 39.123 | Suvorov et al. 2022 |
| Pair 2 | *ananassae* subgroup | *cdtB* | 45 | *setifemur* subgroup | 4 | 24.0026 | This study, time calibrated tree |
| Pair 3 | *saltans* subgroup | *cdtB* | 10 | *cordata/elliptica* subgroups | 8 | 8.2272 | Suvorov et al. 2022 |
| Pair 4 | *cardini* group | *cdtB* | 25 | *guarani* group | 32 | 13.336 | Suvorov et al. 2022 |
| Pair 5 | *Leucophenga* | *cdtB* | 329 | *Amiota/Phortica/Cacoxenus* | 482 | 49 | Dias et al. 2025 |
| Pair 6 | *Rhinoleucophenga* | *cdtB* | 35 | *Gitona* | 22 | 39.3 | Dias et al. 2025 |
| Pair 1 | *melanogaster* 'oriental' clade | PPO2 (PPO3) | 102 | *montium* subgroup | 126 | 22.173 | Suvorov et al. 2022 |
| Pair 2 | *histrio* group | PPO2 | 24 | *immigrans* group | 123 | 21.8046 | This study |
| Pair 3 | *saltans* + *willistoni* groups | PPO2 (PPO4) | 74 | *Lordiphosa* | 99 | 31.297 | Suvorov et al. 2022 |
| Pair 4 | Subfamily Steganinae | PPO2 | 1485 | Subfamily Drosophilinae | 5024 | 67.3 | Dias et al. 2025 |
| Pair 5 | *testacea* group | PPO1 | 8 | *quinaria* group | 45 | 16.6204 | This study |

**Table S7.** PGLS data for effect of *cdtB* presence on diversification rates at three distinct extinction rates.

| **Model** | **Effect of *cdtB* presence on net diversification rate** | **P-Value** | **Fraction of variation explained (Multiple R-squared)** | **Fraction of variation explained (Adjusted R-squared)** |
| --- | --- | --- | --- | --- |
| Net diversification rate (e = 0) ~ cdtb_presence | 0.026079 | 0.307 | 0.104 | 0.0142 |
| Net diversification rate (e = 0.5) ~ cdtb_presence | 0.02426 | 0.233 | 0.139 | 0.0526 |
| Net diversification rate (e = 0.9) ~ cdtb_presence | 0.016756 | 0.205 | 0.156 | 0.0710 |

**Table S8.** Studies surveyed for anti-parasitoid defense mechanisms in the Drosophilidae, related to Figure 1A. See Supplementary File S1 for more details on defense mechanisms for individual species.

| Study | Species surveyed |
| --- | --- |
| Streams 1968 (8) | *D. affinis*, *D. algonquin*, *D. athabasca*, *D. busckii*, *D. immigrans*, *D. melanogaster*, *D. nigromelanica*, *D. paramelanica*, *D. putrida*, *D. robusta* |
| Nappi & Streams 1970 (9) | *D. euronotus*, *D. nigromelanica*, *D. paramelanica*<br>*Screened but no phenotype reported: *D. melanica*, *D. melanura*, *D. micromelanica* |
| Nappi 1970 (10) | *D. euronotus* |
| Nappi 1973 (11) | *D. algonquin* |
| Carton & Kitano 1981 (12) | *D. erecta*, *D. mauritiana*, *D. melanogaster*, *D. orena*, *D. simulans*, *D. teissieri*, *D. yakuba* |
| Schlenke et al. 2007 (13) | *D. ananassae*, *D. biarmipes*, *D. erecta*, *D. eugracilis*, *D. immigrans*, *D. lutescens*, *D. mauritiana*, *D. melanogaster*, *D. paralutea*, *D. pseudoobscura*, *D. robusta*, *D. santomea*, *D. sechellia*, *D. simulans*, *D. teissieri*, *D. virilis*, *D. willistoni*, *D. yakuba*, *Zaprionus vittiger* |
| Havard et al. 2009 (14) | *D. affinis*, *D. azteca*, *D. bifasciata*, *D. guanche*, *D. melanogaster*, *D. miranda*, *D. obscura*, *D. persimilis*, *D. pseudoobscura*, *D. tolteca* |
| Kacsoh 2012 (15) | *D. affinis*, *D. ananassae*, *D. biarmipes*, *D. elegans*, *D. erecta*, *D. eugracilis*, *D. ficusphila*, *D. funebris*, *D. hydei*, *D. immigrans*, *D. kikkawai*, *D. lutescens*, *D. melanogaster*, *D. mojaviensis*, *D. paramelanica*, *D. pseudoobscura*, *D. sechellia*, *D. simulans*, *D. subobscura*, *D. suzukii*, *D. tsacasi*, *D. virilis*, *D. willistoni*, *D. yakuba*, *Zaprionus indianus* |
| Kacsoh et al. 2014 (16) | *Zaprionus indianus*<br>*Hemocytes observed but no infection: *D. affinis*, *D. ananassae*, *D. biarmipes*, *D. elegans*, *D. erecta*, *D. eugracilis*, *D. falleni*, *D. ficusphila*, *D. funebris*, *D. hydei*, *D. immigrans*, *D. kikkawai*, *D. lutescens*, *D. melanogaster*, *D. mimica*, *D. mojaviensis*, *D. palustris*, *D. paramelanica*, *D. phalerata*, *D. pseudoobscura*, *D. putrida*, *D. sechellia*, *D. simulans*, *D. subobscura*, *D. suzukii*, *D. tsacasi*, *D. virilis*, *D. willistoni*, *D. yakuba*, *Scaptodrosophila lebanonensis* |
| Markus et al. 2015 (17) | *D. ananassae*, *D. atripex*, *D. bipectinata*, *D. malerkotiana*, *D. pallidosa*, *D. parabiplectinata*, *D. pseudoananassae*,<br>*Screened and no phenotype reported: *D. auraria*, *D. biarmipes*, *D. bicornuta*, *D. elegans*, *D. erecta*, *D. eugracilis*, *D. ficusphila*, *D. fuyamai*, *D. lucipennis*, *D. lutescens*, *D. mauritiana*, *D. mayri*, *D. melanogaster*, *D. mimetica*, *D. mojaviensis*, *D. nikananu*, *D. persimilis*, *D. prostipennis*, *D. pseudoobscura*, *D. pseudotakahashii*, *D. rhopaloa*, *D. sechellia*, *D. serrata*, *D. teissieri*, *D. simulans*, *D. virilis*, *D. vulcana*, *D. willistoni*, *D. yakuba* |
| Lynch et al. 2016 (18) | *D. erecta*, *D. orena*, *D. mauritiana*, *D. melanogaster*, *D. santomea*, *D. sechellia*, *D. simulans*, *D. suzukii*, *D. teissieri*, *D. yakuba* |
| Iacovone et al. 2018 (19) | *D. melanogaster*, *D. suzukii* |

**Table S9.** Input data for PGLS. Diversification rates were calculated using the <u>bd.ms</u> function in geiger for three extinction rates (0, no extinction; 0.5, moderate extinction; 0.9, high extinction). cdtb_species indicates presence (1) or absence (0) of *cdtB* at the tip. cdtb_prop indicates the proportion of *cdtB*-positive taxa in each clade the tip represents.

| tip | bd.ms_0 | bd.ms_0.5 | bd.ms_0.9 | cdtb_species | cdtb_prop |
| --- | --- | --- | --- | --- | --- |
| GITONA_DISTIGMA | 0.08559149 | 0.067629 | 0.03132872 | 0 | 0 |
| DROSOPHILA_BUSCKII | 0.1195963 | 0.1020939 | 0.06273096 | 0 | 0 |
| DROSOPHILA_NEOCORDATA | 0.252752 | 0.1828177 | 0.06449682 | 0 | 0 |
| RHINOLEUCOPHENGHA_AMERICANA | 0.09844819 | 0.08003488 | 0.04102588 | 1 | 0.5 |
| LEUCOPHENGHA_VARIA | 0.1182869 | 0.104203 | 0.07184614 | 1 | 0.8 |
| AMIOTA_MINOR | 0.1260805 | 0.1119769 | 0.07946651 | 0 | 0 |
| DROSOPHILA_SALTANS | 0.2798747 | 0.2072088 | 0.07801608 | 1 | 0.833 |
| DROSOPHILA_SETIFEMUR | 0.05775601 | 0.03817464 | 0.01093066 | 0 | 0 |
| DROSOPHILA_ANANASSAE | 0.1585938 | 0.1306314 | 0.07025901 | 1 | 0.9375 |
| DROSOPHILA_SUBBADIA | 0.2600849 | 0.2103772 | 0.105887 | 0 | 0 |
| DROSOPHILA_DUNNI | 0.2415594 | 0.1924857 | 0.0918378 | 1 | 0.875 |
| DROSOPHILA_PRIMAEVA | 0.20558 | 0.1878479 | 0.1467224 | 1 | 0.285 |

**Supplementary File S1.** Defense type, genomic resources, and annotation detail data used in this study. Table 1: Defense mechanisms deduced from literature search. Table 2: Genomic resources used in this study. Table 3: Accessions for BLAST queries used in this study. Table 4: Per-species *PPO* counts used for Figure 2D. Table 5: *PPO* annotation details. Table 6: *cdtB* annotation details. Table 7: README for *PPO* and *cdtB* annotation details tables.

